# Learning quality scores for chromatin accessibility bigWig tracks using Machine Learning

**DOI:** 10.64898/2026.06.05.730303

**Authors:** Edward Sanders, Simone G. Riva, Jim R. Hughes

**Affiliations:** MRC Weatherall Institute of Molecular Medicine, Oxford University, Oxford, UK

**Keywords:** ATAC-seq, bigWig, quality control, Machine Learning

## Abstract

High-throughput chromatin accessibility assays such as bulk and single-cell ATAC-seq have generated large collections of processed signal tracks in bigWig format, which are widely used for visualisation, data integration, and Machine Learning (ML)–based analyses. Despite their central role, systematic quality control (QC) frameworks operating directly at the level of bigWig signal tracks remain underdeveloped. This gap limits the ability to assess data reliability and hampers robust downstream analyses.

Here, we present a biologically grounded QC framework for chromatin accessibility bigWig files that integrates peak-level information, background noise estimation, and recovery of stable genomic reference features. Using an ML-based peak caller (LanceOtron), we derive complementary quality metrics capturing signal structure and signal-to-noise properties. We further define constant promoter and CTCF regions as internal biological controls and show that their recovery provides a sensitive measure of data quality across diverse cellular contexts.

We apply this framework to a collection of 502 human chromatin accessibility bigWig tracks spanning a wide range of tissues and cell types. The proposed metrics capture related but non-redundant aspects of signal quality and motivate the use of constant promoter and CTCF recovery as biologically meaningful targets. An XGBoost model trained on LanceOtron-derived features accurately predicts recovery of these stable genomic elements on held-out data (*R*^2^ = 0.97), yielding a continuous and interpretable quality score. Feature importance analysis using SHAP values highlights that model decisions are driven by biologically relevant signal properties rather than arbitrary heuristics. Quantile-based stratification of the quality score is further supported by clear qualitative differences in genome browser visualisations.

Together, this work provides a principled and extensible framework for assessing the quality of chromatin accessibility bigWig tracks, enabling more reliable data integration and supporting downstream ML applications in regulatory genomics.

## I. Introduction

The rapid growth of high-throughput sequencing technologies has led to an unprecedented increase in the scale and diversity of biological datasets, enabling comprehensive exploration of genome function and regulation. Public repositories such as ENCODE and GEO now host vast collections of chromatin accessibility profiles across cell types, tissues, and conditions, providing valuable resources for integrative and comparative analyses [1], [2]. However, this exponential increase in data volume brings substantial challenges in ensuring consistency, reliability, and reproducibility, underscoring the importance of robust quality control (QC) frameworks tailored to specific assay types.

Ensuring data quality is a critical challenge in the analysis of chromatin accessibility assays such as DNase-seq, bulk ATAC-seq, and single-cell ATAC-seq (scATAC-seq). These technologies provide powerful insights into regulatory landscapes, but their outputs are highly sensitive to experimental noise, sequencing biases, and biological variability. DNase-seq, whilst well established, often suffers from artefacts related to sequence preference and digestion bias [3]. Bulk ATAC-seq improves efficiency and resolution but can be affected by PCR amplification biases, transposase sequence preferences, and batch effects that obscure true biological signals [4], [5]. scATAC-seq introduces additional complexity, as its sparse and noisy single-cell profiles complicate peak calling, reproducibility, and the identification of genuine regulatory features [6], [7]. Robust QC pipelines are therefore essential to distinguish technical artefacts from meaningful biological variation and to ensure reliable downstream analyses such as differential accessibility testing, regulatory element discovery, and integrative multi-omics studies. In addition, the increasing use of machine learning (ML) and artificial intelligence (AI) methods in genomics research further amplifies the importance of stringent QC. ML models are highly sensitive to the quality of their training data, and poor input can lead to biased, misleading, or irreproducible results, a notion that unreliable input leads to unreliable output [8], [9]. High-quality, well-curated DNase-seq, bulk ATAC-seq, and scATAC-seq datasets not only enhance the accuracy of predictive models but also improve their generalisability across biological contexts. Conversely, unfiltered noise and technical artefacts can confound feature learning, inflate false positives, and ultimately hinder the discovery of true biological mechanisms [10], [11]. Thus, robust quality assessment is a prerequisite not only for traditional bioinformatics pipelines but also for the reliable application of ML approaches in regulatory genomics.

Whilst QC is well established at the raw read level, particularly with tools such as FastQC for sequencing reads [12], equivalent frameworks for downstream signal tracks such as bigWig files, which represent the final, experiment-level chromatin accessibility signal used for interpretation and downstream analysis, remain underdeveloped. Existing QC approaches primarily focus on read-level metrics or assay-specific heuristics, and do not directly assess processed signal tracks such as bigWigs. BigWig files are widely used to store processed coverage tracks for chromatin accessibility assays, offering efficient visualisation and downstream analysis. However, despite their central role in representing accessibility landscapes, systematic QC approaches for bigWigs are not yet standard practice in the field. This gap leaves researchers without robust methods to evaluate the integrity, reproducibility, or technical biases present at this level of data representation.

In this work, we present a QC framework specifically designed to assess the quality of chromatin accessibility experiments through their bigWig signal tracks, with a focus on bulk and single-cell ATAC-seq data, which encode the final processed representation of experimental accessibility signal. DNase-seq is not explicitly considered, as it shares core experimental principles and systematic biases with ATAC-seq and can therefore be evaluated using closely related QC strategies [4], [13]. By operating directly at the level of processed signal tracks, our approach addresses a notable gap in current state-of-the-art pipelines, enabling systematic assessment of bigWig quality and supporting more robust downstream analyses, large-scale data integration, and ML-based applications.

A central component of our framework is the use of LanceOtron, a ML-based peak caller trained to operate directly on ATAC-seq bigWig files [14]. In many modern analysis workflows, downstream ML or AI models are trained either on the peaks and associated metadata produced by such tools or directly on the underlying bigWig signal itself. In both cases, model performance is intrinsically coupled to the quality of the input signal. Consequently, difficulties encountered by LanceOtron in confidently identifying and characterising peaks provide a practical and biologically informed proxy for the challenges that downstream ML models would face, making its output a natural and effective basis for bigWig-level quality assessment. Importantly, LanceOtron is used here as a feature extraction tool rather than as an object of evaluation, enabling the derivation of signal descriptors that reflect peak structure and confidence. As such, limitations observed at the level of peak calling primarily reflect properties of the underlying experimental signal, rather than artefacts introduced by downstream modelling.

Formally, we define the quality of a chromatin accessibility experiment as its ability to recover biologically stable genomic features from its processed signal. Under this definition, QC becomes a supervised learning problem in which signal-derived features predict recovery of reference elements such as constant promoters and CTCF sites, enabling a principled assessment of experimental signal quality directly from bigWig tracks. In contrast to existing approaches, which typically rely on heuristic or read-level QC metrics, our framework (i) operates directly on processed signal tracks, (ii) defines quality in terms of recovery of biologically stable reference features, and (iii) learns a continuous and interpretable quality score using a supervised machine learning formulation.

## II. Data

To support the development of our QC framework, we assembled a comprehensive collection of chromatin accessibility datasets and integrated them with stable genomic reference features. The data component of this study is organised into two parts: first, the compilation and processing of open chromatin profiles across a wide range of human cell and tissue types, and second, the incorporation of reference genomic features, such as constant promoters and CTCF sites, that serve as internal baselines for evaluating data quality. The following subsections describe these components in detail.

### A. Open chromatin collection

We compiled a collection of 502 distinct cell and tissue types mapped to the human genome assembly GRCh38 (hg38), providing coverage across nearly the entire human body. Specifically, the dataset includes: 10 cell types from [15]; 221 cell types from [16], equally divided between adult and fetal cells; 23 cell types from [17]; 19 primary peripheral blood mononuclear cell (PBMC) types (see Table I); 175 immune cell types from [18]; 8 erythropoiesis cell types from [19]; 12 pancreatic PBMC samples from [20]; 15 erythropoiesis samples from [21]; 4 human embryonic stem cell (hESC) samples from GEO accession GSE124853; and 15 immune cell samples from GEO accession GSE125164.

**TABLE I:** Human Healthy Donors, 10X Genomics, accessed 04/03/2023.

Dataset Name with hyperref URL
|  |
| --- |
| 500 PBMC (Next GEM v1.1) |
| 1k PBMC (Next GEM v1.1) |
| 5k PBMC (Next GEM v1.1) |
| 10k PBMC (Next GEM v1.1) |
| 10k PBMC, ATAC v1.1, Chromium X |
| 10k PBMC, ATAC v1.1, Chromium Controller |
| 3k PBMC - Granulocytes Removed Through Cell Sorting |
| 10k PBMC - Granulocytes Removed Through Cell Sorting |
| 3k PBMC - No Cell Sorting |
| 10k PBMC - No Cell Sorting |
| 10k PBMCs, Multiome v1.0, Chromium X |
| 10k PBMCs, Multiome v1.0, Chromium Controller |

Datasets from [15], [17], and PBMCs were analysed in greater detail, both upstream and downstream, to isolate and generate bigWig files corresponding to the relevant cell types of interest. The data from [16] were obtained directly in bigWig format from their interactive web atlas. In contrast, datasets from [18], [19], [20], [21], as well as GEO accessions GSE124853 and GSE125164, were re-processed using the CATCH-UP pipeline [22]. In total, all 502 bigWig tracks were subjected to peak calling with LanceOtron [14] to identify significant regions of open chromatin for downstream enrichment analyses. The collections are summarised in Table II.

**TABLE II:** Summary of collected datasets. The table reports the dataset names, the number of distinct cell types represented in each, and their corresponding source or publications. Each dataset provides a unique set of chromatin accessibility profiles derived from diverse cellular contexts.

| Collection name | Number different cell types | Authors |
| --- | --- | --- |
| CAD | 10 | [15] |
| catlas_fetal | 111 | [16] |
| catlas_adult | 110 | [16] |
| MPAL | 23 | [17] |
| super_PBMC | 19 | 10x Genomics |
| Calderon | 175 | [18] |
| Ludwig | 8 | [19] |
| pancreatic_PBMC | 12 | [20] |
| erythropoiesis | 15 | [21] |
| hESC | 4 | GSE124853 |
| immune_cell | 15 | GSE125164 |

### B. Reference Genomic Features for QC

In addition to the introduced epigenetic data, we incorporated reference genomic features to establish stable baselines for QC. Gene annotations were obtained from RefSeq (NCBI catalog GCF 000001405.40) and used to define constant promoter regions, i.e., promoter sequences associated with housekeeping genes that are consistently active across diverse cell types and conditions. These regions serve as internal positive controls, since their accessibility is expected to remain stable and largely independent of cell-type-specific regulatory programs. As such, robust recovery of promoter accessibility provides a sensitive indicator of data quality, coverage, and signal-to-noise ratio, rather than biological variability. To derive a set of constant genes, we employed a previously published list of housekeeping genes [23], yielding 3,688 genes. Given the objectives of our analysis, we prioritised conservatism over completeness in order to ensure robustness, minimising the inclusion of contextdependent or weakly expressed promoters that could confound quality assessment.

Similarly, to define constant CCCTC-binding factor (CTCF) sites-genome-wide binding regions of the CTCF protein, a key architectural regulator of 3D genome organisation and chromatin looping [24], we relied on the curated list reported in [25]. Using a conservative selection, we identified 4,636 constant CTCF sites. Unlike promoters, CTCF sites primarily reflect structural rather than transcriptional regulatory activity and are therefore expected to exhibit accessibility across a broad range of cellular contexts. Their consistent detection provides an orthogonal control to promoter-based metrics, enabling assessment of data quality across both transcriptionally active and architectural elements of the genome. Together, constant promoters and CTCF sites provide complementary and biologically grounded reference features for evaluating the integrity and reliability of chromatin accessibility bigWig tracks.

To further clarify, constant promoters typically produce strong and robust accessibility signals, reflecting high and consistent transcriptional activity across cell types, whereas constant CTCF sites tend to exhibit weaker and more spatially constrained accessibility patterns associated with architectural binding. Considering both feature classes jointly allows the QC framework to assess experimental signal quality across a range of signal strengths, ensuring sensitivity to both high-signal and more challenging, lower-signal regulatory elements.

## III. Method

To systematically evaluate the quality of bigWig tracks, we designed a framework composed of complementary metrics that capture different aspects of chromatin accessibility data integrity. Each component of the framework targets a specific source of variability or artefact, and together they provide a robust basis for assessing dataset reliability. An overview of the full workflow is shown in Figure 1. In the following subsections, we describe: (A) peak calling, which assesses the consistency of identified accessible regions and provides complementary statistics; (B) background noise, which quantifies unwanted signal levels; (C) constant promoters, used as a baseline for accessibility across datasets; (D) constant CTCF regions, serving as additional internal standards; and (E) an integrated XGBoost-based quality assessment, which leverages all preceding metrics to produce a unified and data-driven measure of bigWig quality.

**Figure 1.**
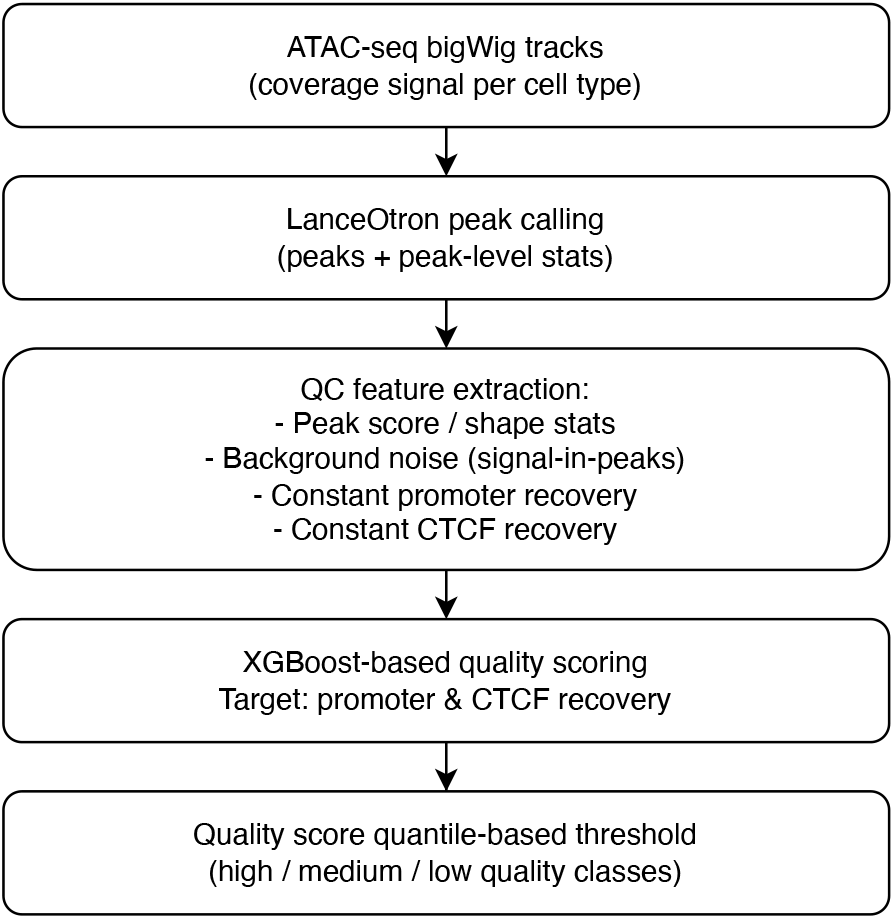
Graphical overview of the bigWig QC framework. ATAC-seq bigWig tracks are peak-called using LanceOtron, from which multiple QC features are extracted, including peak statistics, background noise estimates, and enrichment at constant promoter and CTCF regions. These features are integrated using an XGBoost model trained to predict recovery of stable genomic elements, producing a continuous quality score that is partitioned into quality classes using quantile-based thresholds.

### A. Peak calling

All bigWig tracks were subjected to peak calling using LanceOtron [14], a supervised ML framework designed to identify regions of significant chromatin accessibility. LanceOtron annotates each peak with two key metrics: a peak score, reflecting the overall statistical confidence of the peak, and a shape score, capturing the conformity of the signal profile to an expected accessibility peak shape. In addition to these two primary measures, LanceOtron provides a rich set of auxiliary statistics describing peak characteristics and local signal properties, which together offer a detailed representation of peak quality and signal structure.

From these annotations, we derived straightforward QC metrics, including the mean peak score and the mean shape score across all peaks within a track, which serve as intuitive indicators of data quality. Well-behaved bigWig tracks typically yield high values for both metrics, as high-quality data allow LanceOtron to confidently and consistently identify well-defined peaks. Conversely, noisy or low-quality tracks often result in reduced peak and shape scores, reflecting uncertainty in peak detection and diminished signal-to-noise separation. Beyond these summary statistics, the full set of LanceOtron-derived features was incorporated as input to the downstream XGBoost model (see Section III-E), enabling a more comprehensive, multivariate assessment of bigWig quality.

We selected LanceOtron over traditional peak callers such as MACS2 and its successor MACS3 [26] because it integrates deep learning with heuristic scoring, providing not only sensitive and accurate peak detection but also quality-related metrics that are particularly informative for assessing bigWig track reliability. Moreover, LanceOtron operates directly on bigWig files, eliminating the need to revert to raw read formats (e.g., BAM) and thereby streamlining the QC workflow. Crucially, because LanceOtron itself is a deep learning model trained to recognise biologically meaningful accessibility patterns, its ability, or failure, to confidently characterise peaks provides a direct and sensitive proxy for the suitability of a bigWig track for downstream ML and AI applications.

### B. Background noise

To quantify background noise, we defined a metric that measures the proportion of signal attributable to annotated peak regions relative to the total signal in each bigWig track. In this context, background noise refers to signal that is broadly distributed across the genome and not associated with discrete, biologically meaningful regions of chromatin accessibility, arising from technical artefacts, sequencing biases, or non-specific transposition events. By contrast, true accessibility signal is expected to be concentrated within well-defined peak regions. Signal quantification was performed using the `pyBigWigstats` method (with `exact=True`) to obtain precise coverage values. Total signal was computed across all autosomal chromosomes to avoid confounding effects arising from sex chromosome copy number differences. To further characterise signal-to-noise properties, this calculation was repeated for peaks selected at increasing LanceOtron peak score thresholds. Comparing the proportion of signal retained at more stringent cutoffs provides a measure of how effectively a dataset concentrates signal within high-confidence accessible regions. High-quality bigWig tracks are expected to retain a substantial fraction of signal within strong peaks, even at stringent thresholds (e.g., peak score > 0.95). In contrast, noisy or low-quality tracks typically exhibit a larger proportion of signal outside peak regions, indicative of elevated background noise and a reduced signal-to-noise ratio.

### C. Constant promoters

To assess the stability of promoter accessibility across datasets, we relied on a previously published list of constant housekeeping genes [23]. For each gene, transcription start site (TSS) coordinates were obtained from RefSeq annotations (NCBI GCF 000001405.40). Promoter regions were defined by extending a 100 bp window upstream and downstream of each TSS, as informed by observations using REgulamentary [27]. This window size effectively captures promoter-associated accessibility while minimising overlap with enhancers or architectural elements such as CTCF sites. Enhancers were intentionally excluded because their accessibility is often highly cell-type- and condition-specific, making them unsuitable as stable reference regions for QC [28], [29]. The resulting definition, therefore, provides a robust and conservative set of constant promoter regions.

For each dataset, we quantified the proportion of constant promoters that overlap at least one called peak, the proportion that overlap a high-confidence peak, and the fraction of the total signal localised within these promoter regions. High-quality bigWig tracks are expected to exhibit consistent and reproducible accessibility at constant promoters, reflected in high overlap rates and a substantial concentration of signal. In contrast, datasets with poor coverage or elevated background noise tend to show reduced promoter recovery, indicating limited sensitivity in detecting universally accessible regulatory regions. Because housekeeping gene promoters are expected to remain accessible across cell types and conditions, their consistent detection provides a biologically grounded internal control that enables separation of technical data quality effects from genuine biological variability.

### D. Constant CTCF

In addition to promoter regions, we evaluated dataset quality using constant CTCF sites. CTCF (CCCTC-binding factor) is a ubiquitously expressed DNA-binding protein that plays a central role in 3D genome architecture and chromatin looping [24]. We used a curated list of constant CTCF binding sites [25], representing genomic loci where CTCF occupancy is consistently observed across a wide range of cell types and conditions.

For each dataset, we quantified the proportion of constant CTCF sites that overlap at least one called peak as well as the proportion that overlap a high-confidence peak. High-quality datasets are expected to show strong and reproducible accessibility at these sites, reflected in high overlap rates even at stringent peak score thresholds. In contrast, noisy or low-quality datasets typically show reduced CTCF site recovery, indicating limited sensitivity or elevated background noise.

Because constant CTCF sites correspond to stable architectural elements of the genome rather than transcriptionally regulated features, their consistent detection provides an orthogonal and biologically grounded internal control for quality assessment. Incorporating CTCF-based metrics alongside promoter-based measures enables evaluation of data quality across both regulatory and structural dimensions of chromatin accessibility, strengthening the robustness of the overall QC framework.

### E. XGBoost-based quality scoring

A high-quality bigWig track is expected to exhibit strong and consistent recovery of constant promoters and constant CTCF sites whilst maintaining low background noise, reflecting reliable experimental signal across both high-amplitude regulatory regions and weaker, architecturally defined binding sites. To formalise this intuition, we trained an XGBoost model [30] in which the target variables were the recovery rates of constant promoter and constant CTCF regions per track. For clarity, this corresponds to the number of constant promoter and CTCF regions recovered per track, proportional to recovery rate given a fixed reference set. XGBoost was selected because it efficiently captures non-linear relationships and higher-order interactions among heterogeneous input features, whilst remaining robust to feature collinearity and varying feature scales. In this formulation, bigWig quality is learned directly from biologically grounded reference features, rather than from subjective or manually defined quality labels.

The input features to the model were derived primarily from LanceOtron peak calling. LanceOtron provides a rich set of peak-level annotations and summary statistics that capture both the confidence and structural properties of accessible regions, including peak score, shape score, peak abundance, and signal distribution characteristics. These LanceOtron-derived features were combined with background noise measurements and enrichment metrics at constant promoter and constant CTCF regions to form the final feature set. Together, these inputs allow the model to learn how properties of peak structure and signal distribution relate to the reliable recovery of stable genomic elements.

Model training and hyperparameter selection were performed using a multi-stage grid search strategy, enabling progressive refinement of tree depth, learning rate, and regularisation parameters to balance predictive performance and model stability. The final XGBoost model represents the best-performing configuration identified during this optimisation procedure and was selected based on its ability to accurately predict constant promoter and CTCF recovery across the full collection of bigWig tracks (see Table III).

**TABLE III:** Final XGBoost hyperparameters selected via multi-stage grid search. Model selection was performed using five-fold cross-validation with negative mean squared error as the optimisation metric.

| Hyperparameter | Selected value |
| --- | --- |
| learning_rate | 0.1 |
| n_estimators | 100 |
| max_depth | 4 |
| min_child_weight | 2 |
| subsample | 0.6 |
| colsample_bytree | 1.0 |
| gamma | 0 |
| reg_alpha | 0.1 |
| reg_lambda | 0.5 |
| cv | 5 |
| scoring | neg_mean_squared_error |

The trained model outputs a continuous quality score for each bigWig track, reflecting its predicted ability to recover constant promoter and CTCF regions. To categorise tracks into quality classes, thresholds were defined using quantiles of the score distribution, which were visually assessed using boxplot representations in the Results section. This strategy enables data-driven separation of high- and low-quality tracks based on the empirical distribution of model scores, rather than relying on fixed or arbitrary cutoffs.

By grounding quality thresholds in quantile-based partitions of the learned score distribution, this approach preserves interpretability whilst remaining flexible across heterogeneous datasets. Importantly, the resulting quality classification is directly anchored to biologically meaningful sensitivity in recovering stable regulatory and architectural elements of the genome, providing a robust and extensible framework for bigWig QC.

We note that the objective of this work is not to benchmark alternative predictive models, but to establish a biologically grounded formulation of bigWig-level quality assessment. As such, the focus is on demonstrating that the proposed feature set captures meaningful signal properties and enables accurate prediction of recovery of stable genomic elements, rather than on comparing model performance across different learning algorithms. While simple baselines such as linear models based on peak counts or coverage could be considered, these do not capture the multi-dimensional structure of signal features exploited by the proposed framework.

## IV. Results

In this section, we present the results of applying the proposed QC framework to a large collection of chromatin accessibility experiments, evaluated through their corresponding bigWig signal tracks. We first examine the behaviour of individual QC metrics, focusing on background noise, recovery of constant promoter regions, and recovery of constant CTCF sites (Subsection IV-A), in order to characterise how data quality varies across datasets and how these metrics relate to one another. We then evaluate the performance of the XGBoost-based quality scoring model, assessing its ability to predict recovery of stable genomic features and using model interpretability analyses to identify which features most strongly contribute to quality discrimination (Subsection IV-B). Finally, we present genome browser visualisations [31] of representative high-, intermediate-, and low-quality tracks selected based on quantiles of the model-derived quality score, providing qualitative validation of the scoring framework through direct inspection of chromatin accessibility signal (Subsection IV-C).

Together, these results illustrate both the empirical properties of the proposed QC metrics and the effectiveness of their integration into a unified, data-driven quality assessment.

### A. Behaviour of QC metrics across bigWig tracks

To characterise the variability in data quality across the collected bigWig tracks, we first examined the behaviour of the individual QC metrics derived from peak calling, background noise estimation, and recovery of constant genomic features. Together, these metrics provide complementary views of signal integrity, coverage, and sensitivity to stable regulatory elements. Importantly, they capture complementary aspects of experimental signal quality, reflecting both global noise characteristics and sensitivity to biologically stable regulatory features.

Background noise, quantified as the fraction of signal not attributable to called peaks, showed substantial variation across datasets (Figure 2). Tracks with low background noise exhibited a higher concentration of signal within high-confidence peaks, whereas tracks with elevated background noise displayed broadly distributed signal, consistent with reduced signal-to-noise ratio. When stratifying tracks by background noise levels, we observed a clear degradation in the recovery of constant genomic features as noise increased.

**Figure 2.**
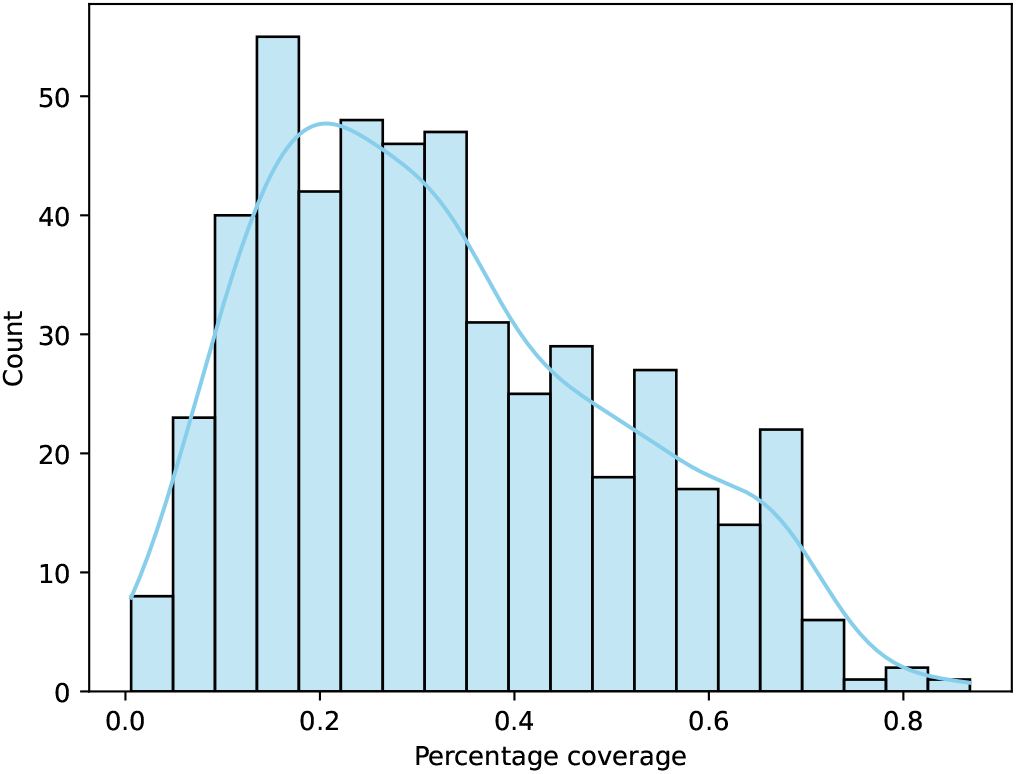
Distribution of the percentage of signal covered by called peaks across all bigWig tracks. This metric reflects background noise levels, with lower values indicating a larger fraction of signal distributed outside peak regions.

Consistent with this trend, recovery of constant promoter regions decreased progressively with increasing background noise (Figure 3). Tracks with low background noise recovered a large fraction of housekeeping gene promoters, with accessibility profiles that were both strong and spatially well-defined. In contrast, tracks with high background noise showed markedly reduced promoter recovery, indicating diminished sensitivity to universally accessible regulatory regions.

**Figure 3.**
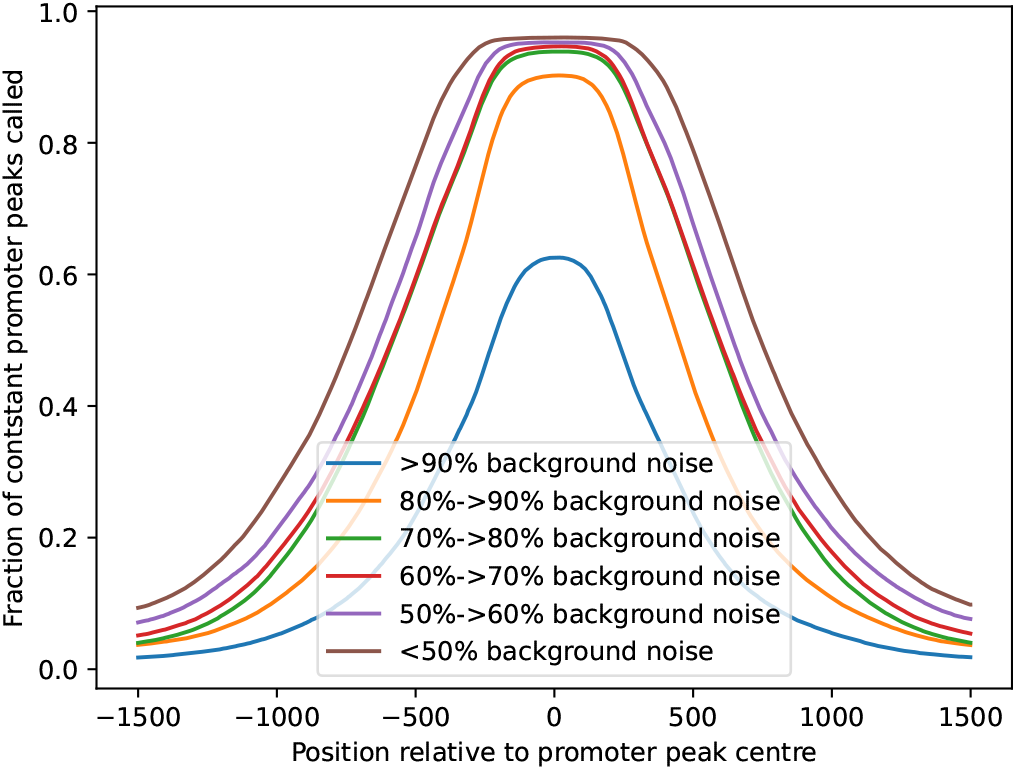
Positional recovery of constant promoter regions across bigWig tracks stratified by background noise levels. The fraction of constant promoter peaks detected is shown relative to the promoter peak centre. Tracks with lower background noise exhibit higher and more sharply localised promoter recovery, whereas increasing background noise leads to reduced and more diffuse recovery profiles.

A similar pattern was observed for constant CTCF sites (Figure 4). Because CTCF binding sites represent stable architectural elements rather than transcriptionally regulated regions, their recovery provides an orthogonal measure of data quality. High-quality tracks consistently recovered a large proportion of constant CTCF sites, whilst tracks with increasing background noise showed a pronounced loss of detectable CTCF-associated accessibility. Notably, CTCF recovery appeared particularly sensitive to elevated background noise, reinforcing its utility as a stringent internal control for chromatin accessibility data.

**Figure 4.**
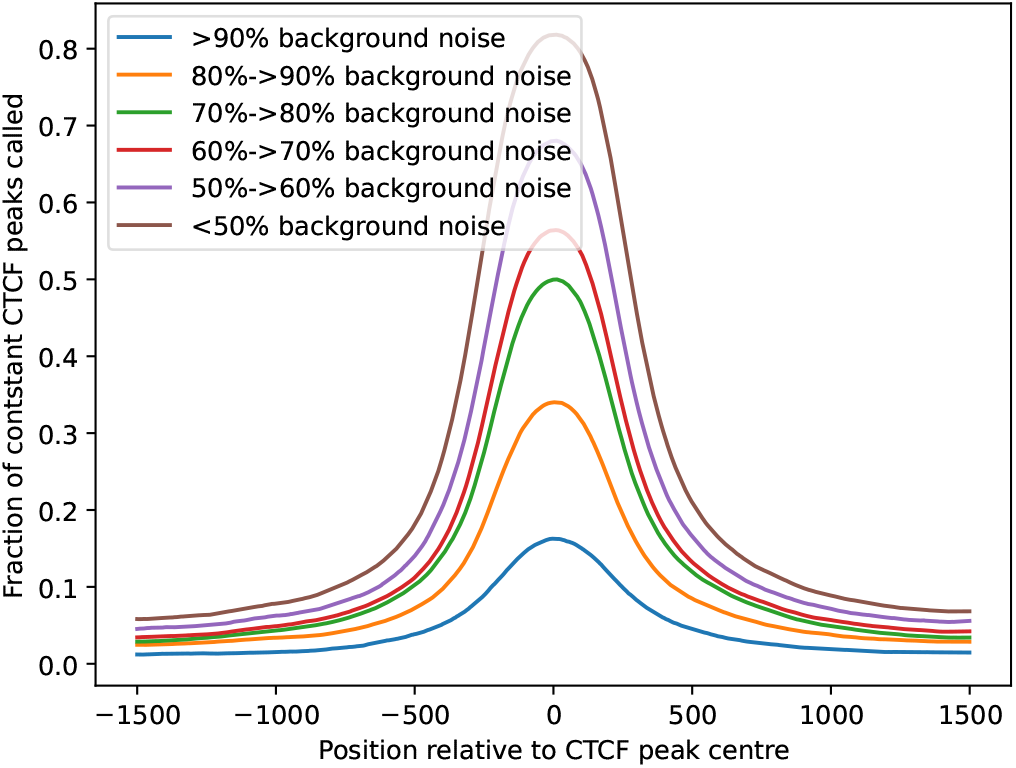
Positional recovery of constant CTCF sites across bigWig tracks stratified by background noise levels. The fraction of constant CTCF peaks detected is shown relative to the CTCF peak centre. Tracks with lower background noise exhibit higher and more sharply localised CTCF recovery, whereas increasing background noise results in reduced and broadened recovery profiles.

This difference in behaviour reflects the distinct signal properties of the two feature classes: constant promoters generally yield strong, high-amplitude accessibility signals, whilst constant CTCF sites represent weaker and more spatially restricted signals. As a result, successful recovery of both promoters and CTCF sites indicates that an experiment exhibits sufficient signal-to-noise characteristics across a spectrum of signal strengths, providing a more stringent and comprehensive assessment of experimental signal quality than either feature alone.

Taken together, these results demonstrate that background noise, constant promoter recovery, and constant CTCF recovery capture related but non-redundant aspects of bigWig quality. Whilst background noise reflects global signal contamination, promoter and CTCF recovery directly measure the ability of a dataset to detect biologically stable regulatory and architectural features. Importantly, this combination of metrics captures precisely the biological signal of interest for downstream analyses, making recovery of constant promoter and CTCF regions a natural and biologically grounded target for training the quality scoring model.

### B. XGBoost-based quality scoring and feature importance analysis

Building on the observed relationships among QC metrics, we evaluated the performance of the XGBoost model trained to predict recovery of constant promoter and CTCF regions from LanceOtron-derived features and background noise measurements. Following the optimisation procedure described in Section III-E, the dataset was randomly split such that 80% of the bigWig tracks were used for model training, whilst the remaining 20% were held out for independent testing. Model selection and hyperparameter tuning were performed exclusively on the training set.

On the held-out test set, the model showed strong agreement between predicted and observed counts of constant promoter and CTCF peaks across bigWig tracks (Figure 5). Quantitatively, model performance achieved a mean squared error (MSE) [32] of 86,294.05 and a coefficient of determination (*R*^2^) [33] of 0.9735, indicating that the model explains the vast majority of variance in recovery of stable genomic features. These results demonstrate that the selected input features are highly informative of biologically meaningful accessibility signals and that the learned quality score provides an accurate and robust proxy for the underlying experiment’s ability to recover constant promoter and CTCF regions from its chromatin accessibility signal.

**Figure 5.**
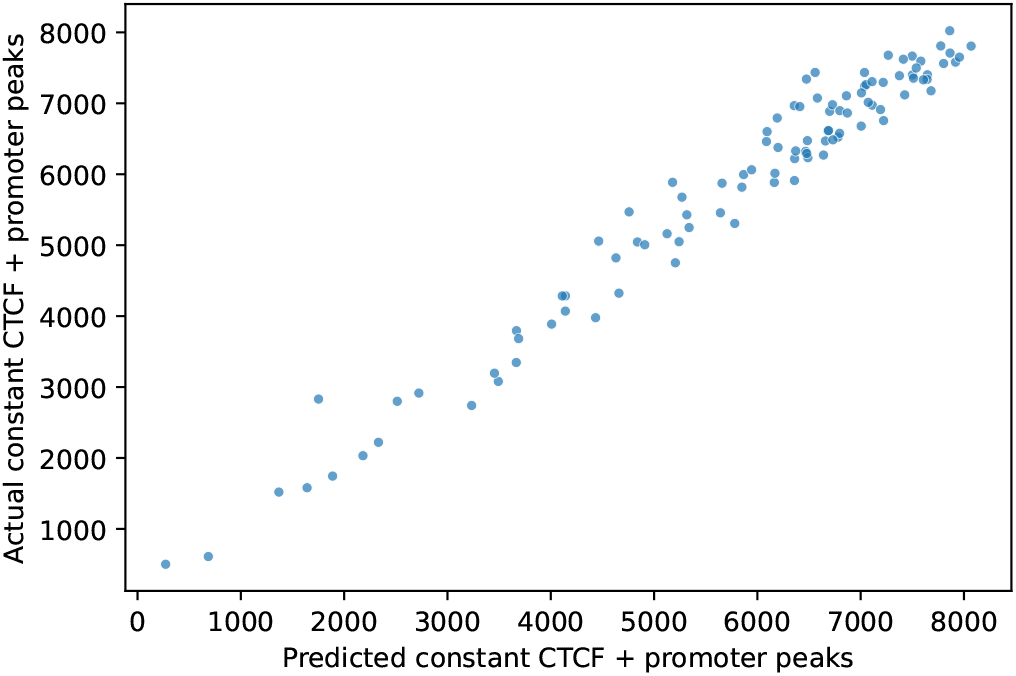
Comparison of predicted and observed counts of constant promoter and CTCF peaks on the held-out test set. Each point represents a bigWig track. The strong agreement along the diagonal indicates accurate prediction of recovery of stable genomic features by the XGBoost model.

To better understand how individual features contributed to the model’s predictions, we analysed feature importance using SHAP (SHapley Additive exPlanations) values [34] (Figure 6). This analysis revealed that the total number of peaks, enrichment-related metrics, and measures of signal concentration within peaks were among the most influential features driving model output. In particular, peak abundance and enrichment scores exhibited strong positive contributions to predicted promoter and CTCF recovery, whilst features associated with elevated background noise contributed negatively.

**Figure 6.**
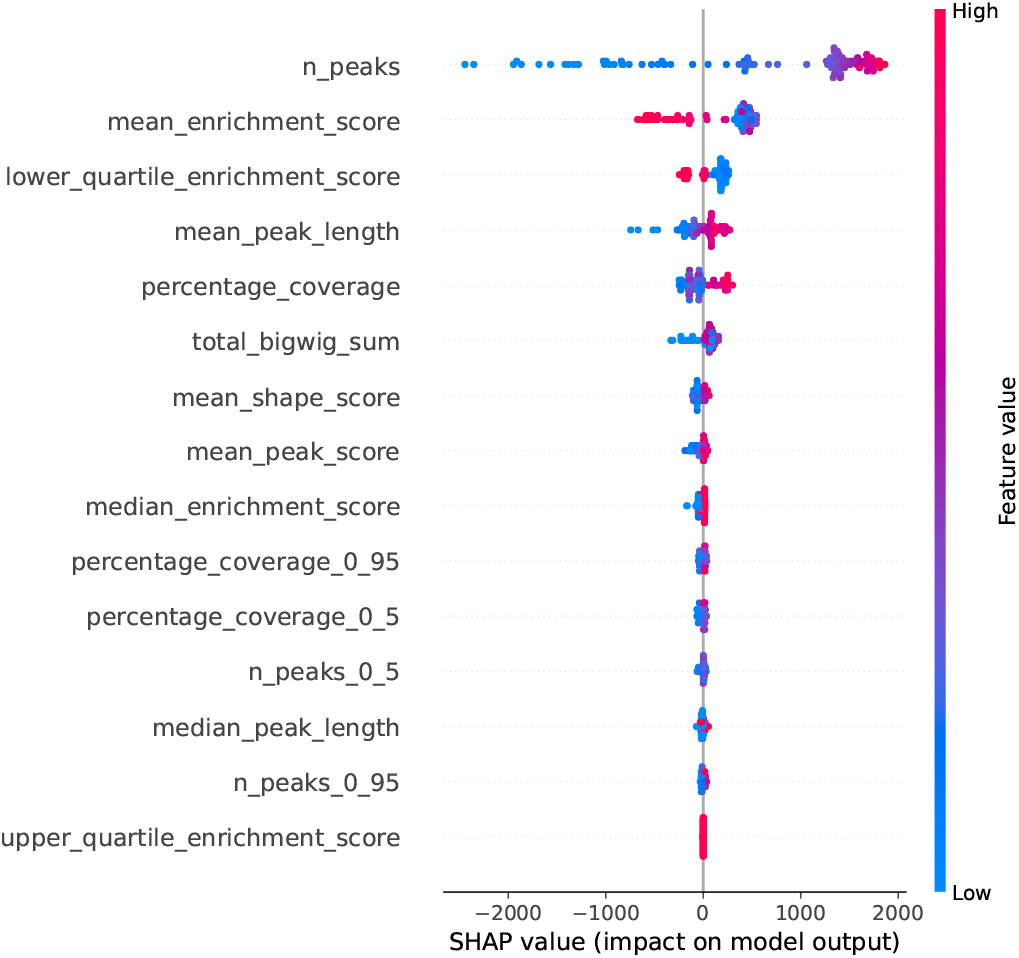
SHAP summary plot showing the contribution of individual QC features to the XGBoost model output. Each point represents a bigWig track, coloured by feature value. Features are ordered by their overall impact on model predictions, highlighting the relative importance of peak abundance, enrichment metrics, and signal coverage in discriminating track quality.

Importantly, SHAP analysis highlighted that no single metric was sufficient to explain model behaviour in isolation. Instead, the model relied on combinations of peak-level statistics, signal distribution measures, and enrichment features to discriminate between high- and low-quality tracks. This finding supports the use of an integrated, multivariate approach to quality assessment, rather than reliance on individual heuristics or fixed thresholds.

Overall, the XGBoost-based quality score provides a quantitative and interpretable summary of bigWig quality that is directly grounded in biologically meaningful recovery of constant genomic features. By combining multiple complementary QC metrics and leveraging model interpretability via SHAP values, this approach enables robust discrimination of dataset quality whilst offering insight into the factors that drive model decisions. These results indicate that recovery of constant promoter and CTCF regions can be accurately inferred from signal-derived features alone, supporting the validity of the proposed formulation of experimental signal quality.

### C. Visual inspection of representative high- and low-quality tracks

To complement the quantitative evaluation of model performance, we next examined representative examples of the underlying bigWig signal in genomic context. Using the quantile-based quality score distribution shown in the boxplot representation (Figure 7), we stratified bigWig tracks into high-, intermediate-, and low-quality groups. From each quantile, we randomly selected a small number of tracks from the held-out test set and visualised their chromatin accessibility signal using a genome browser (Figure 8). These differences reflect variation in experimental signal quality rather than biological differences at the locus shown, which is held constant across tracks.

**Figure 7.**
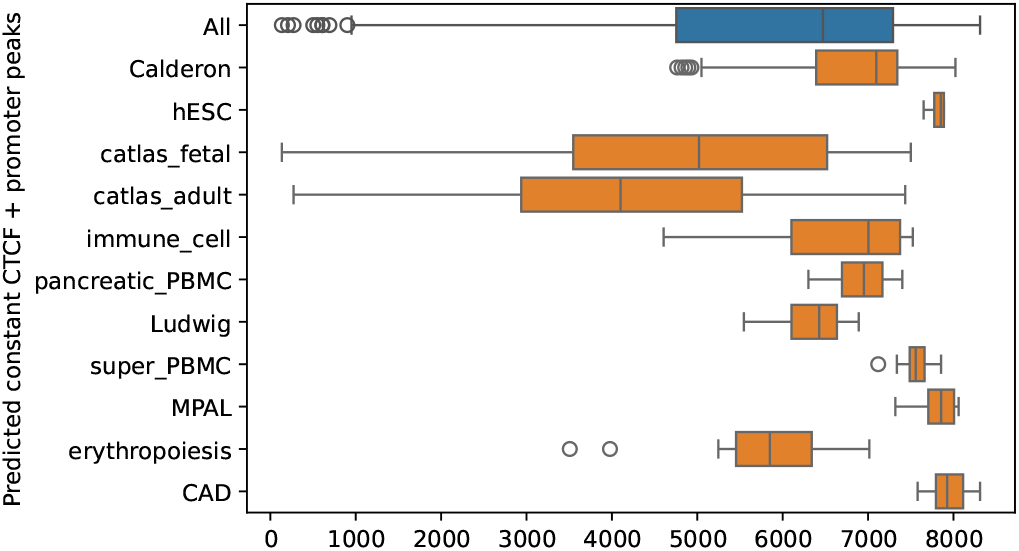
Distribution of the XGBoost-predicted recovery of constant promoter and CTCF regions across all bigWig tracks and stratified by dataset. Boxplots summarise the distribution of predicted recovery values within each collection, highlighting variability in signal quality across different chromatin accessibility datasets.

**Figure 8.**
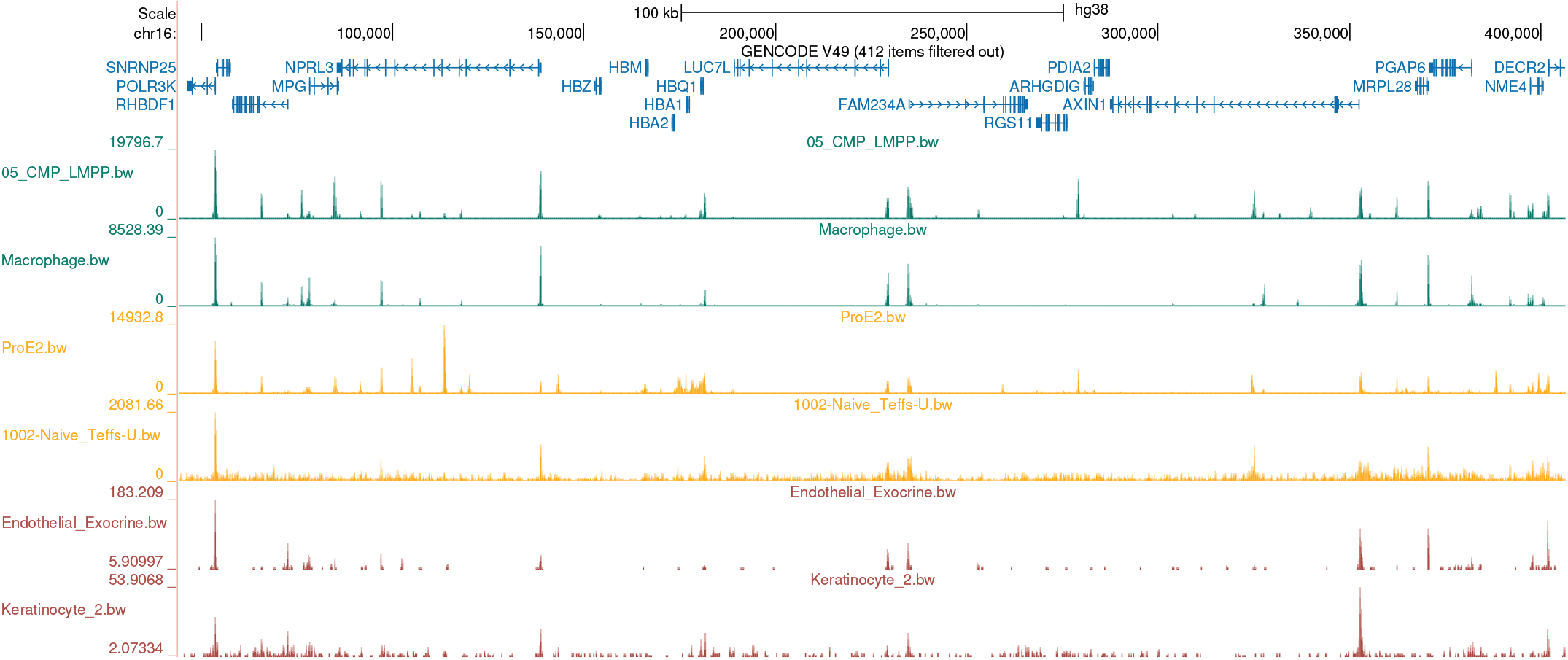
Genome browser visualisation of representative bigWig tracks stratified by quantiles of the XGBoost-derived quality score. Tracks shown in green correspond to high-quality bigWigs (upper quantile), tracks shown in orange correspond to intermediate-quality bigWigs (above or below the median of the score distribution), and tracks shown in red correspond to low-quality bigWigs (lower quantile). All tracks are displayed over the same genomic locus to illustrate systematic differences in peak definition, background noise, and recovery of constant promoter and CTCF regions.

These genome browser views reveal clear qualitative differences that are consistent with the model-derived quality scores. High-quality tracks exhibit sharp, well-defined peaks with low background signal and clear enrichment at constant promoter and CTCF regions. Tracks from intermediate quantiles show broader peaks and moderate background noise, whilst low-quality tracks display diffuse signal with poor peak definition and limited recovery of stable genomic features. Importantly, these examples demonstrate that the quantile-based quality score reflects not only abstract model outputs but also interpretable differences in raw accessibility signal, providing intuitive validation of the proposed QC framework.

## V. Conclusion & Future Work

In this work, we introduced a QC framework for assessing the quality of chromatin accessibility experiments through their bigWig signal tracks that operates directly at the level of processed signal data. By combining peak-level information from an ML-based peak caller with complementary metrics capturing background noise and recovery of constant promoter and CTCF regions, which together capture experimental signal quality across a range of signal strengths, our approach provides a biologically grounded assessment of data quality beyond traditional read-level QC. The analysis of over 500 bigWig tracks spanning a wide range of cell and tissue types demonstrates that these metrics capture related but non-redundant aspects of signal integrity and sensitivity to stable regulatory and architectural elements of the genome.

We further showed that these QC features can be effectively integrated using an XGBoost model trained to predict recovery of constant promoter and CTCF regions, yielding a continuous and interpretable quality score. Importantly, this approach evaluates the integrity of the experimental signal itself, rather than properties of individual analysis tools or downstream models. The strong agreement between predicted and observed recovery on held-out data, together with feature importance analysis using SHAP values, highlights that model decisions are driven by biologically meaningful properties of the signal rather than arbitrary heuristics. Importantly, quantile-based stratification of the quality score was reflected in clear and intuitive differences in genome browser visualisations, providing qualitative validation that the proposed framework captures interpretable differences in chromatin accessibility signal.

Overall, this work addresses a critical gap in current chromatin accessibility analysis pipelines by providing a principled and extensible approach for assessing the quality of bigWig signal tracks. By anchoring quality assessment to recovery of stable genomic features, the proposed framework supports more reliable downstream analyses, large-scale data integration, and the application of ML models in regulatory genomics.

Several directions for future work arise naturally from this study. First, whilst the present framework focuses on ATAC-seq–derived bigWig tracks, the underlying principles are broadly applicable to other chromatin accessibility assays, including DNase-seq and emerging multi-omics protocols. Extending and benchmarking the approach across additional assay types and species will further strengthen its general applicability.

Second, future work could explore incorporating additional reference features, such as constitutive enhancers or other structural elements, to refine quality assessment in specific biological contexts. Similarly, integrating temporal or replicate-level information may enable more nuanced discrimination between technical artefacts and genuine biological variability.

Finally, the quality score developed here could be directly integrated into downstream analysis pipelines, for example, by informing sample filtering, weighting tracks in large-scale integrative studies, or serving as an explicit input to ML models trained on chromatin accessibility data. As ML-driven approaches continue to expand in genomics, robust and interpretable QC at the level of processed experimental signal tracks will be increasingly essential.

## Acknowledgment & Declaration

S.G.R. is supported by the MRC grant (MC UU 00029/3). E.S. is supported by the Wellcome Trust grant (225220/Z/22/Z). J.R.H. is supported by the Wellcome Trust grants (225220/Z/22/Z and 106130/Z/14/Z) and the MRC grant (MC UU 00029/3).

J.R.H. is a co-founder and director of Nucleome Therapeutics and provides consultancy to the company.

